# Role of two metacaspases in development and pathogenicity of the Rice Blast fungus, *Magnaporthe oryzae*

**DOI:** 10.1101/2020.12.10.420794

**Authors:** Jessie Fernandez, Victor Lopez, Lisa Kinch, Mariel A. Pfeifer, Hillery Gray, Nalleli Garcia, Nick V. Grishin, Chang-Hyun Khang, Kim Orth

## Abstract

Rice blast disease caused by *Magnaporthe oryzae* is a devastating disease of cultivated rice worldwide. Infections by this fungus lead to a significant reduction in rice yields and threats to food security. To gain better insight into growth and cell death in *M. oryzae* during infection, we characterized two predicted *M. oryzae* metacaspase proteins, MoMca1 and MoMca2. These proteins appear to be functionally redundant and are able to complement the yeast Yca1 homologue. Biochemical analysis revealed that *M. oryzae* metacaspases exhibited Ca^2+^ dependent caspase activity *in vitro*. Deletion of both *MoMca1* and *MoMca2* in *M. oryzae* resulted in reduced sporulation, delay in conidial germination and attenuation of disease severity. In addition, the double Δ*Momca1mca2* mutant strain showed increased radial growth in the presence of oxidative stress. Interestingly, the Δ*Momca1mca2* strain showed an increase accumulation of insoluble aggregates compared to the wild-type strain during vegetative growth. Our findings suggest that MoMca1 and MoMca2 promote the clearance of insoluble aggregates in *M. oryzae*, demonstrating the important role these metacaspases have in fungal protein homeostasis. Furthermore, these metacaspase proteins may play additional roles, like in regulating stress responses, that would help maintain the fitness of fungal cells required for host infection.

**IMPORTANCE:** *Magnaporthe oryzae* causes rice blast disease that threatens global food security by resulting in the severe loss of rice production every year. A tightly regulated life cycle allows *M. oryzae* to disarm the host plant immune system during its biotrophic stage before triggering plant cell death in its necrotrophic stage. The ways *M. oryzae* navigates its complex life cycle remains unclear. This work characterizes two metacaspase proteins with peptidase activity in *M. oryzae* that are shown to be involved in the regulation of fungal growth and development prior to infection by potentially helping maintain fungal fitness. This study provides new insight into the role of metacaspase proteins in filamentous fungi by illustrating the delays in *M. oryzae* morphogenesis in the absence of these proteins. Understanding the mechanisms by which *M. oryzae* morphology and development promote its devastating pathogenicity may lead to the emergence of proper methods for disease control.

## INTRODUCTION

Caspases are a family of conserved cysteine-dependent, aspartate-specific proteases that play an essential role in metazoan programmed cell death. They regulate multiple cellular behaviors that contribute to organism fitness and pathology (1). This family of proteases contains a unique α/β hemoglobinase fold that consists of a large subunit (p20) containing the catalytic histidine/cysteine dyad and small subunit (p10) (2). Caspases are synthesized as zymogens, and, upon an external stimulus, are activated by autocatalysis. At present, there are no known caspase homologs in non-metazoan organisms; however, a group of distantly-related orthologous caspases known as metacaspases was discovered in fungi, plant and protozoa (2–5). Metacaspases share structural homology to components of animal caspases, but lack substrate specificity for aspartate residues (3). Instead, they specifically cleave substrates preceded by positively-charged lysine and arginine residues in the P1 position (3, 5). Bioinformatics analysis identified three types of metacaspases based on the presence or absence of a N-terminal prodomain with the caspase domain organization (p20/p10 subunits) (6). For instance, budding yeast *S. cerevisiae* harbors a single type I metacaspase known as Yca1 (4). Similar to animal caspases, Yca1 contains the conserved Cys-His catalytic dyad and undergoes autoproteolytic processing to yield p20 (~20 kDa) and p10 (~12 kDa) subunits from the inactive zymogen (4). The N-terminal prodomain is rich on poly-Q/N repeats, a motif predicted to be involved in self-aggregation (7).

Overall, metacaspases are multifunctional proteases essential for normal physiology of non-metazoan groups. For instance, the loss or inactivation of Yca1 in yeast alters the timing of the cell cycle progression by elongating the G1 phase and perturbing the G2/M mitotic checkpoint, implicating Yca1 in the regulation of the cell cycle (8). Moreover, Yca1 stimulates apoptotic-like cell death during oxidative stress and aging in yeast (4, 9–12). In the absence of this caspase, yeast cells tolerate low doses of H_2_O_2_ promoting their survival under these harsh conditions (4). Yca1 has also been implicated in the clearance of insoluble protein aggregates, thereby protecting cells against byproducts of aging and toxic amyloids (7). Studies demonstrated that Yca1 interacts with known members of the proteostasis network such as Cdc48, Hsp104, and the Hsp70/40 chaperone systems (7). Indeed, the loss of Yca1 results in the accumulation of stress response chaperones leading to the enrichment of autophagic bodies as a compensatory response of survival (7). Further investigations established that Yca1 maintains proteostasis through direct interaction with the ubiquitin proteasome system (13). Specifically, the ubiquitination of Yca1 was shown to have a direct impact on its function within the proteostasis network; once Yca1 was ubiquitinated, the yeast was able to regulate protein aggregation levels and autophagy (13). Metacaspases have also been discovered in a few other fungi groups like *Aspergillus spp.*, *Podospora anserina*, *Candida albicans* and recently in the corn smut, *Ustilago maydis* (14–18). Like Yca1 in yeast, most of these fungal caspases have been linked to stress and age-related responses as consequences of several environmental stimuli (14–18).

The filamentous fungus*, Magnaporthe oryzae,* is the causative agent of rice blast, one of the most destructive diseases of cultivated rice in the world, and is responsible for the annual destruction of approximately 10-30% of the rice harvested globally (19). During infection, *M. oryzae* undergoes extensive developmental changes which allow it to break into plant cells, build elaborate infection structures, and proliferate inside host cells without causing visible disease symptoms (19). The infection starts when the three-celled conidium attaches to the surface of the rice leaf. The conidium germinates and forms a germ tube. The tip of the germ tube differentiates into an infection structure called the appressorium (20, 21). Once the appressorium matures, turgor pressure builds within the structure and drives the protrusion of the penetration peg, enabling it to breach the leaf cuticle (19, 22). Subsequently, *M. oryzae* colonizes the plant tissue, sporulates and spreads by air to uninfected rice plants to continue its life cycle.

Here, we identify and characterize two *M. oryzae* metacaspases, MoMca1 and MoMca2, which exhibit a C14 peptidase activity. While the Cys/His catalytic dyad is essential for its autocatalytic processing, these metacaspases require Ca^2+^ for their full proteolytic activity *in vitro,* suggesting a critical role *in vivo* for this catalytic mechanism. In the absence of both *MoMca1* and *MoMca2*, *M. oryzae* exhibits delayed conidial germination, and subsequent delayed appressorium formation. Moreover, the double mutant strain, but not the single mutant strains, was impaired in developing typical WT lesions on rice leaves compared to the WT strain suggesting that MoMca1 and MoMca2 are redundant and that both are required for full pathogenicity. Consistent with this observation, both enzymes can functionally complement the yeast Δ*yca1* strain. Interestingly, we observed that these proteins promote the clearance of insoluble aggregates to maintain the fitness of *M. oryzae* cells. Collectively, our study demonstrates that the *MoMca* genes play important roles in growth, conidiation, appressorium development, and pathogenesis for *M. oryzae*.

## RESULTS

### *M. oryzae* genome contains two putative metacaspase proteins

To investigate critical roles of metacaspases in *M. oryzae* life cycle, we first searched the *Magnaporthe* genome for metacaspase-related genes. We chose to analyze this group of proteins because they have been extensively studied in yeast, but remain largely uncharacterized in filamentous, plant pathogenic fungi. Using the protein sequence of yeast Yca1 for BLASTP searches, we identified two putative metacaspase genes MGG_04626 and MGG_13530 (now named *MoMca1* and *MoMca2*, respectively) in the *M. oryzae* genome database. *MoMca1* and *MoMca2* genes are located on chromosomes, Chr3 and Chr4, respectively. These putative proteases are predicted to encode a typical type I metacaspase containing an N-terminal prodomain with a Q/N rich repeat motif and a C-terminal peptidase-C14 caspase domain (Fig. 1A). MoMca1 and MoMca2 encode 396 and 410 amino acid proteins, respectively. Clan clustering analysis of 2712 metacaspase protein sequences showed that *M. oryzae* metacaspases clustered in the same clan as yeast Yca1, with their catalytic domains sharing approximately 45% sequence similarity (Fig. 1B). Bioinformatics analysis also revealed MoMca1 and MoMca2 encode 67% similarity between their catalytic domains. Using ClustalW, the sequence alignment of MoMca1 and MoMca2 with Yca1 illustrated that *M. oryzae* proteins contain the predicted histidine and cysteine as the catalytic dyad specific for clan CD proteases (Fig. 1A and C). In figure S1A, the tertiary structure of full-length *M. oryzae* metacaspases is modeled using the structural template of yeast Yca1 according to Swiss-Model (PDB:4f6p) (23).

**Figure 1.**
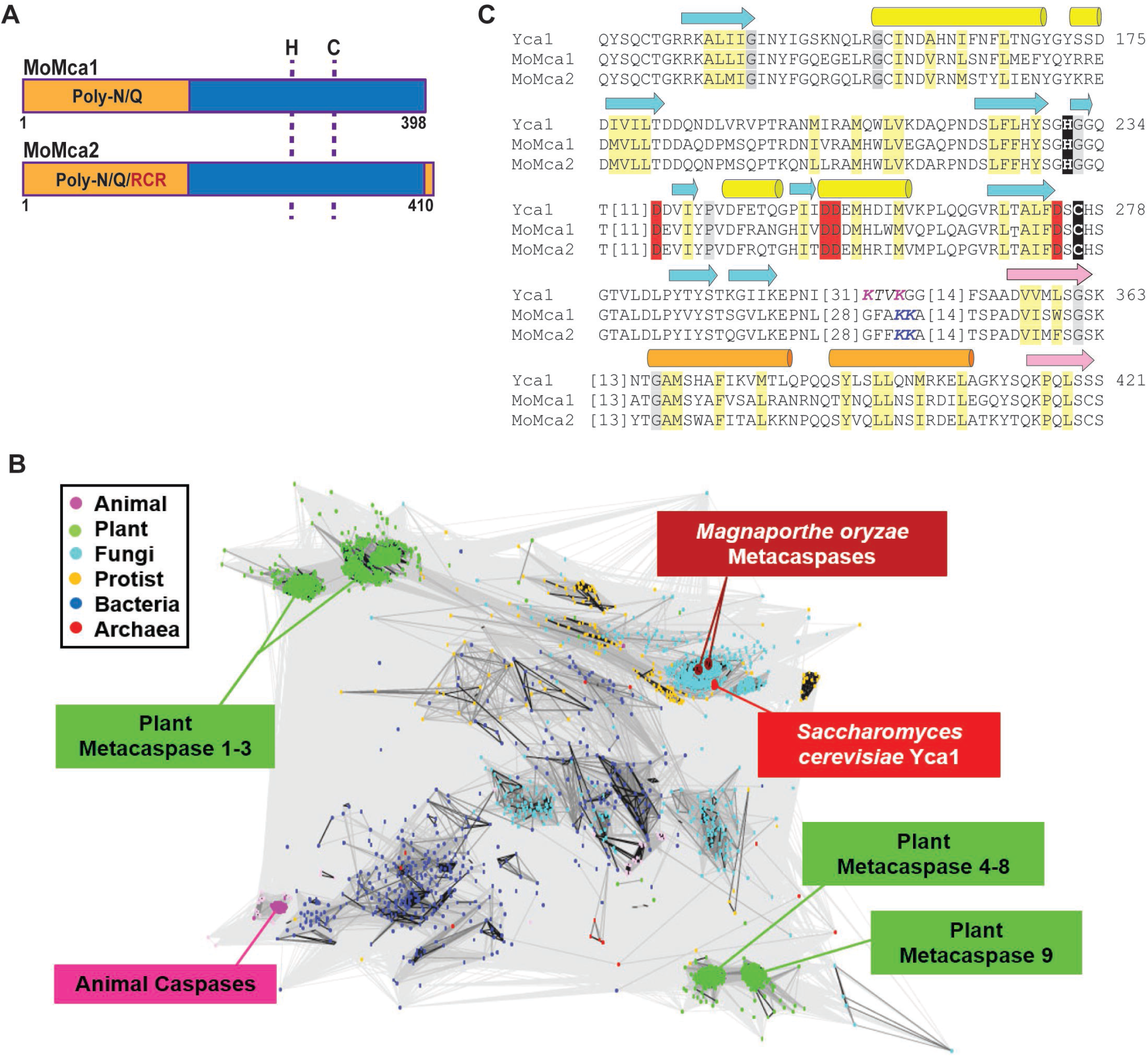
Bioinformatics identification of two *Magnaporthe oryzae* metacaspases. (**A**) Schematic representation of MoMca1 (MGG_04926) and MoMca2 (MGG_13530). Pro-domain, rich in asparagine (N) and glutamine (Q) residues, is indicated in orange. The caspase domain is marked in blue. RCR domain: partial sequence motif of chitin synthesis regulation labeled in red. Dashed lines show the position of the histidine (H) and cysteine (C) catalytic dyad (**B**) CLANS clustering analysis is represented graphically as the network of BLAST-derived sequence similarities between 2712 metacaspase protein sequences. Dots represent sequences, with the shade of connecting lines denoting similarity from gray (low) to black (high). (**C**) The catalytic sites of MoMca1 and MoMca2 are aligned with Yca1, highlighting conserved active site residues (black), calcium binding residues (red), Yca1 cleavage site (magenta), predicted MoMca1/MoMca2 cleavage site (Blue), mainly hydrophobic positions (light yellow), and mainly small positions (gray). Beta-strands (arrow) and alpha/helices (cylinder) from the Yca1 structure are indicated above the alignment and colored cyan/yellow prior to and pink/orange after the cleavage site (italics).

### MoMca1 and MoMca2 are Ca^2+^-dependent proteases

For most of the metacaspases studied so far, autocatalysis of an inactive zymogen is responsible for generating an active enzyme monomer (4, 5, 23). Therefore, we addressed the question whether MoMca1 and MoMca2 undergo autocatalytic activation in a similar manner as yeast Yca1. Due to the insolubility of both *M. oryzae* metacaspase proteins in bacterial cells, we decided to use an *in vitro* transcription/translation approach to enable a rapid expression of the full-length protein of MoMca1 and MoMca2. A recent study demonstrated that *Arabidopsis* metacaspase AtMCP2d and yeast Yca1 strictly require Ca^2+^ for their proteolytic activity *in vitro* (5). We observed that recombinant MoMca1 and MoMca2 are already autocatalytically processed upon translation, yielding a ~35 kDa intermediate common to both metacaspases and the p10 subunit, at ~12 kDa in MoMca1 and ~10 kDa in MoMca2 (Fig. 2A). To examine whether treatment with divalent cations such as Ca^2+^ positively affect *M. oryzae* metacaspases autoprocessing, we added 1mM CaCl_2_ to the translation mixture and observed further autocatalysis yielding a ~25 kDa fragment of the large subunit (Fig. 2A). These results demonstrate that Ca^2+^ specifically enhanced the autocatalytic processing of MoMca1 and MoMca2.

**Figure 2.**
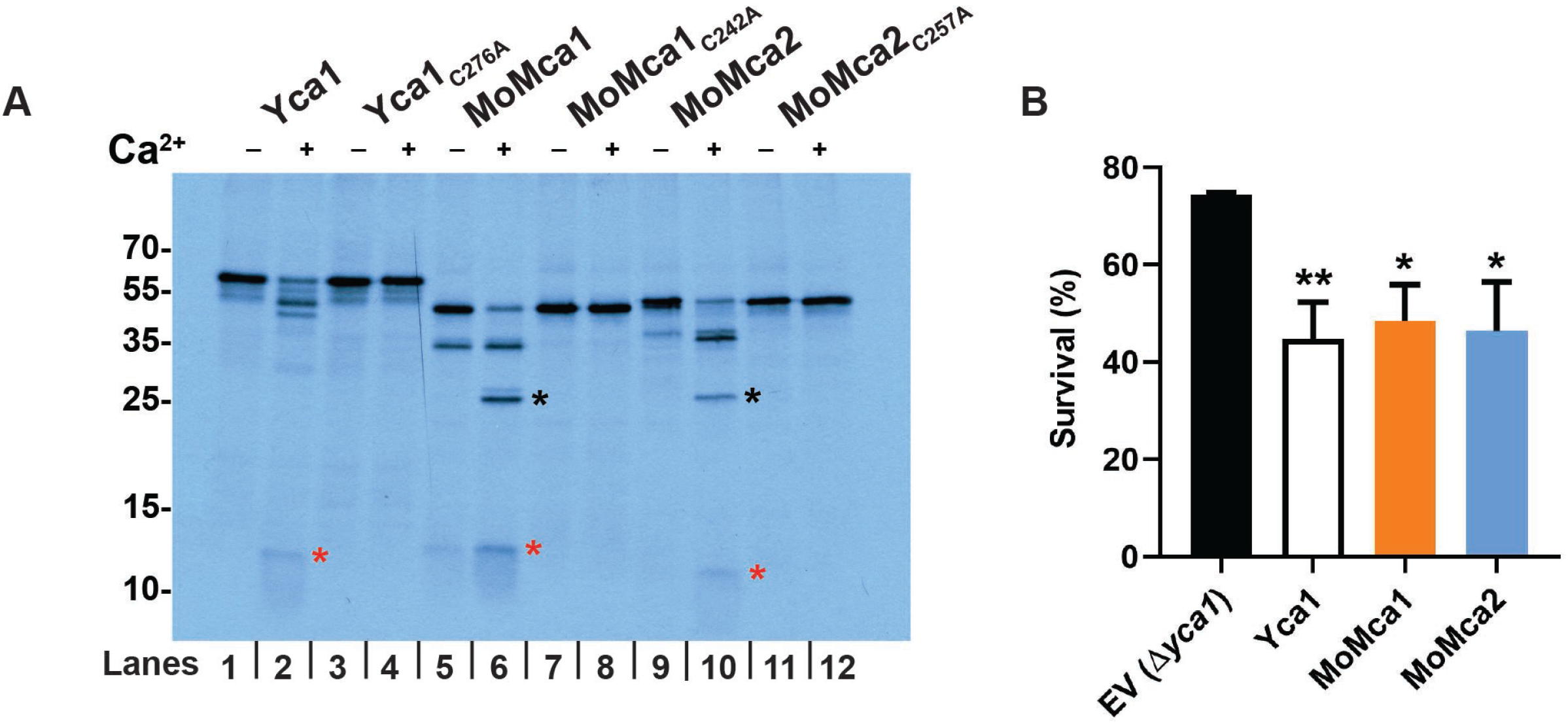
Biochemical characterization of *Magnaporthe oryzae* metacaspases. (**A**) Protease activity assay of MoMca1 and MoMca2 in the presence or absence of 1mM CaCl_2_ *in vitro*. Red and black asterisks indicate the p20 and p10 subunits, respectively. Samples were loaded in pairs; odd lanes are controls for even lanes (**B**) Survival assay of Δ*yca1* cells expressing Yca1, MoMca1, MoMca2, and the corresponding empty vector control (EV Δ*yca1*) after 24 hours of galactose induction shown as % survival in the presence of 1.2mM H_2_O_2_ treatment. Data are represented as the mean of three independent measurements. Error bars denote SD. Asterisks indicate statistically significant differences (*p<0.05, **p<0.01, ***p<0.001, one-way ANOVA with Tukey’s multiple comparisons test using GraphPad Prism 8).

Sequence alignment of MoMca1 and MoMca2 with Yca1 showed the predicted catalytic Cys/His dyad residues in both *M. oryzae* metacaspases (Fig. 1B). The catalytic cysteine (Cys^242^ for MoMca1 and Cys^257^ for MoMca2) was replaced by an alanine residue in their coding sequence. Figure 2A shows that in the presence of Ca^2+^, the catalytic dead proteins are unable to autoprocess and generate catalytic subunits, remaining as inactive zymogens. These results demonstrated that the predicted conserved cysteines are essential for metacaspase autoproteolysis.

### Expression of MoMca1 and MoMca2 in a heterologous system

Previous studies demonstrated that low doses of H_2_O_2_ promote the expression of the Yca1 in yeast, leading to the induction of apoptosis (4). The deletion strain Δ*yca1* exhibits higher rates of survival rate under oxidative stress conditions compared to the wild type strain (4). To determine if MoMca1 and MoMca2 functionally complement Yca1 in H_2_O_2_-mediated apoptosis, we cloned the *M. oryzae* metacaspases into the Δ*yca1* yeast strain and grew these strains in liquid galactose inducing media containing 1.2mM H_2_O_2_ for 24 h. Figure 2B shows a significant reduction in cell survival observed upon complementation of the Δ*yca1* strain containing either *MoMca1* or *MoMca2* compared to the Δ*yca1* strain carrying the empty vector. These results indicate that MoMca1 and MoMca2 can functionally complement Yca1 when expressed in *S. cerevisiae*.

### MoMca1 and MoMca2 are required for the virulence of *M. oryzae*

To determine if *MoMca1* and *MoMca2* play a role in *M. oryzae* pathogenicity, we generated single Δ*Momca1* and Δ*Momca2* deletion strains, a double mutant strain, Δ*Momca1mca2*, and a complemented strain, Δ*Momca1mca2-C*, containing pBGt MoMCA1MCA2 expression vector (Fig. S1B and C). These strains were evaluated for pathogenicity on rice leaves. We inoculated susceptible rice plants with spore suspensions of the wild-type (WT) or the mutant strains. The WT strain exhibited the expected necrotic lesions on rice leaves. By contrast, the Δ*Momca1mca2* strain displayed a significant reduction in disease severity compared to the WT and the single mutants. (Fig. 3A and B). This data suggests that metacaspases are required for full pathogenicity in rice blast fungus, signifying their importance in *M. oryzae* pathogenicity.

**Figure 3.**
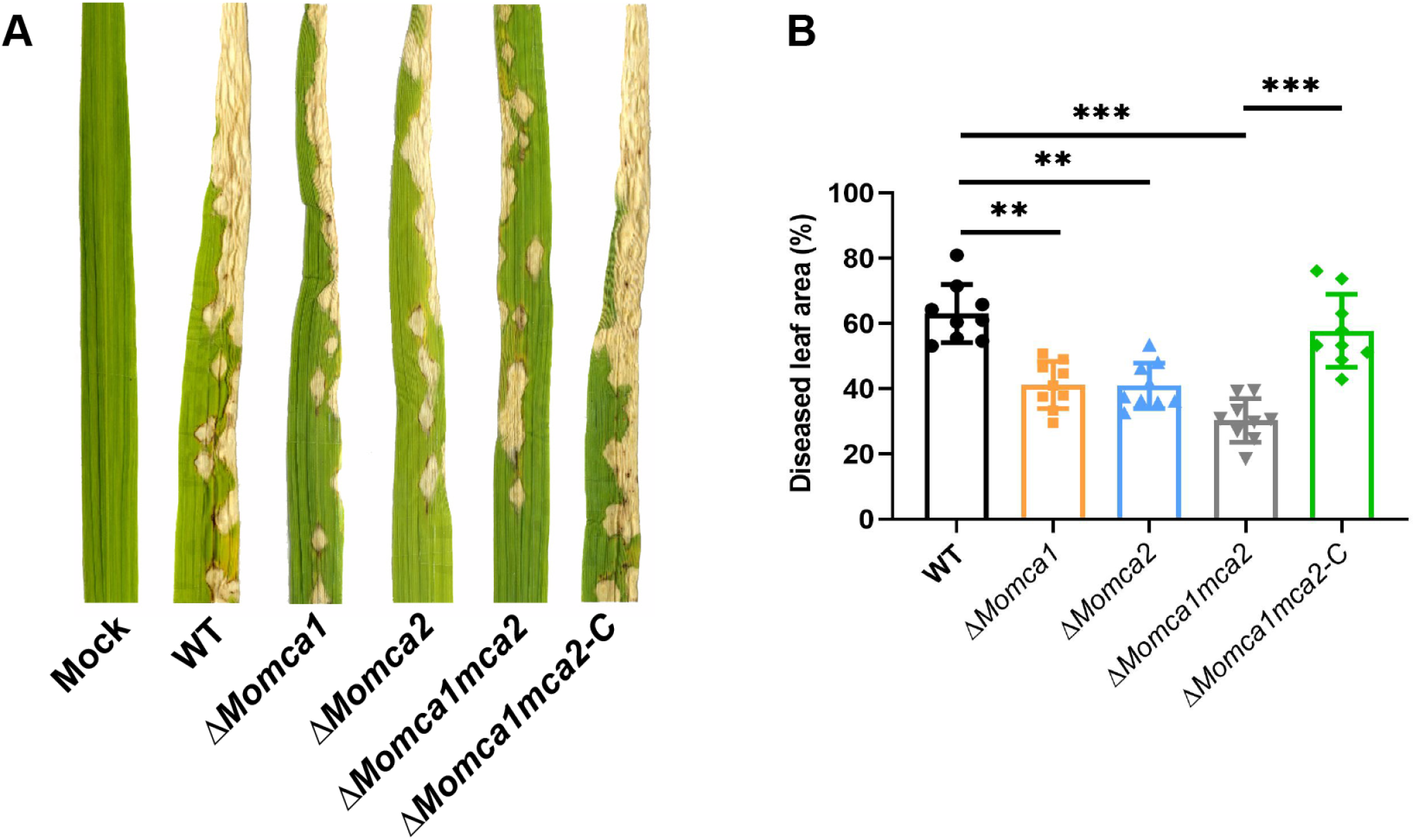
*MoMca1* and *MoMca2* are required for full pathogenicity. (**A**)(**B**) Δ*Momca1mca2* strain was reduced in virulence compared to WT when applied to leaves of the susceptible rice variety YT16. Δ*Momca1mca2*-C strains refers to the complemented strain. (**B**) Diseased leaf area was measured using ImageJ. Results are the mean of three independent measurements. Error bars denote SD. Asterisks indicate statistically significant differences (**p<0.01, ***p<0.001, one-way ANOVA with Tukey’s multiple comparisons test using GraphPad Prism 8).

### *M. oryzae* metacaspases are required for sporulation

To investigate the role of MoMca1 and MoMca2 in growth and development of *M. oryzae*, we assessed the WT, mutant and complemented strains for colony diameter on complete media (CM; Fig. S2A and B). However, the Δ*Momca1mca2* strain displayed severely reduced sporulation when compared to the WT and single mutant strains (Fig. S2C). Moreover, none of these strains showed differences in sensitivity to cell-wall assembly inhibitor or osmotic stress (Fig. S3A and B). Taken together, this data suggests that metacaspases are individually dispensable for spore development and appropriate vegetative growth on axenic conditions, and are redundant in providing an essential function for spore development.

### MoMca1 and MoMca2 are expressed under oxidative conditions *in vivo*

To further verify whether MoMca1 and MoMca2 impact *M. oryzae* survival under oxidative stress as Yca1 does in yeast, we first measured the transcript levels of *MoMca1* and *MoMca2* under different stress conditions. A previous study showed that H_2_O_2_ induces the expression of Yca1 leading to the activation of programmed cell death in yeast (4). Therefore, we used qPCR to analyze the expression of *MoMca1* and *MoMca2* in the WT *M. oryzae* strain under H_2_O_2_ and the free radical generator, menadione. Only *MoMca1* expression was induced in the presence of H_2_O_2_ whereas both metacaspases were induced with menadione, suggesting that the enzymes may play distinct roles in responding to cell stress (Fig. 4A).

**Figure 4.**
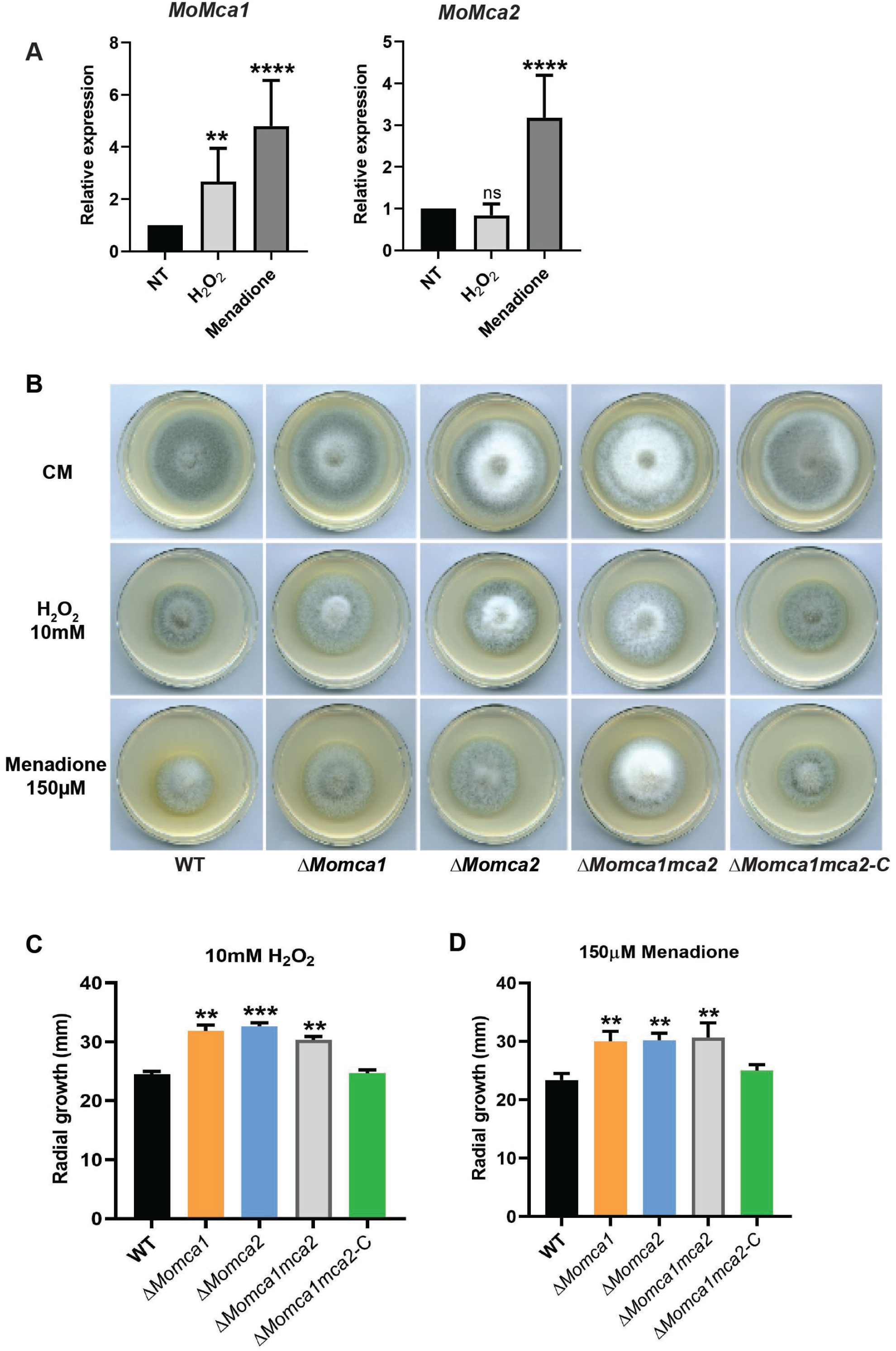
The Δ *Momca1,* Δ *Momca1* and Δ *Momca1* Δ *mca2* mutant strains showed resistance to the oxidative stress conditions compared to the WT strain. (**A**) Quantitative RT-PCR analysis showing the expression of *M. oryzae* metacaspase-encoding genes, *MoMca1* and *MoMca2* in the WT KV1 strain under oxidative stress conditions. Fungal mycelia were grown in liquid CM for 48h at 25°C with agitation (150 rpm) before treatment with either 5mM H_2_O_2_ or 100μM menadione for 1hr. Expression was normalized to the housekeeping gene *ACT1*. Data are 2-ΔΔCq ± SD, N = 3 experiments. NT = no treatment (**B**)(**C**)(**D**) WT KV1 and Δ*Momca* strains were inoculated as 10 mm mycelial plugs onto 55 mm diameter plates of complete media containing H_2_O_2_, or menadione at the concentrations indicated. Images were taken at 7 days after growth. (**C**)(**D**) Measurements of radial growth of *M. oryzae* WT and mutant strains mycelia under oxidative stress conditions. Results are the mean of three independent measurements. Error bars denote SD. ns: not significant. Asterisks indicate statistically significant differences (**p<0.01, ***p<0.001, ****p<0.0001, one-way ANOVA with Tukey’s multiple comparisons test using GraphPad Prism 8).

To measure the sensitivity of the mutants and survival under oxidative stress conditions, we grew the WT and mutant strains on CM plates containing H_2_O_2_ or menadione and compared their radial growth to the WT strain. Interestingly, we observed that both the single and double mutants were less sensitive to H_2_O_2_ and menadione conditions than the WT and complemented strains (Fig. 4B). The mutant strains also exhibited higher radial growth under stress than the WT and complemented strains (Fig. 4C and D). These results indicate that the *M. oryzae* metacaspases play an important role in stress-induced programmed cell death.

### Deletion of *MoMca1* and *MoMca2* leads to a delay in conidial germination and appressorium formation

Next, we addressed whether *MoMca1* and *MoMca2* might play a role in conidial germination or appressorium development. We collected spores from 10-day old CM plates of the WT and mutant strains, and placed spores in suspension on a hydrophobic surface. At 3 and 5 hours post incubation (hpi), the WT strain germinated and began to form immature appressoria (Fig. 5A and B). By contrast, the Δ*Momca1mca2* strain was impaired in germination and appressorium morphogenesis (Fig. 5A-C). The single mutants, Δ*Momca1* and Δ*Momca2*, showed non-significant reduction in germination rates. However, Δ*Momca2* showed a reduction in appressorium development at 3 hpi, but appeared similar to WT at 5 hpi (Fig. 5A-C). At 12 and 20 hpi, all the strains were able to develop appressoria (Fig. 5C). Interestingly, at 20 hpi we observed that approximately 20% of the germinated spores in the Δ*Momca1mca2* strain showed elongated germ tubes compared to those of the WT and single mutants (Fig. 6A and B). This data suggests that *M. oryzae* metacaspases play an important, but redundant, role in conidial germination, as there is a delay in appressorium development in their absence.

**Figure 5.**
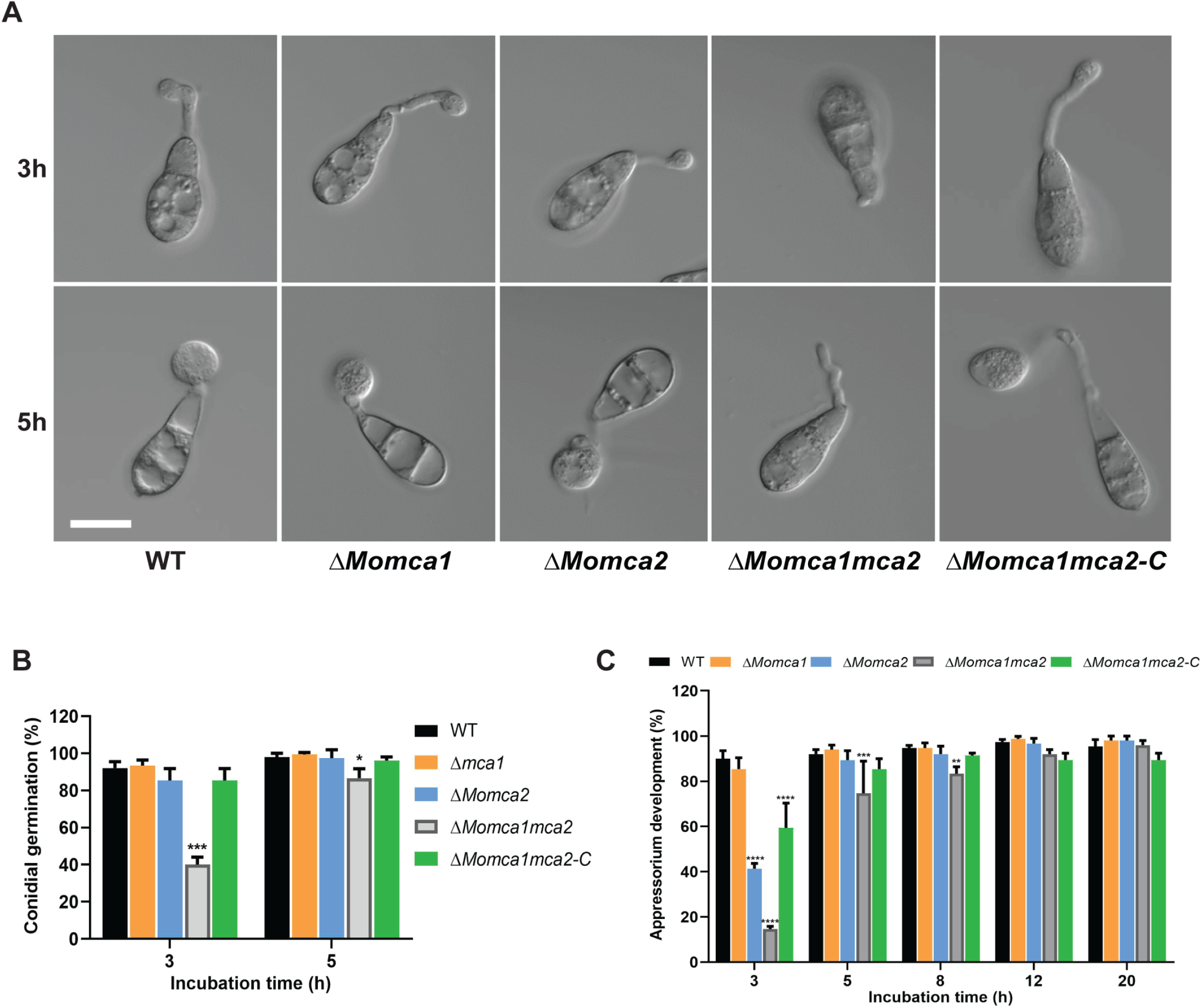
Loss of metacaspase genes delays conidial germination and subsequent formation of appressorium on hydrophobic surfaces. (**A**) Spores of WT KV1, Δ*Momca1,* Δ*Momca2,* Δ*Momca1mca2*, and Δ*Momca1mca2*-C were applied to artificial hydrophobic surfaces (coverslips). (**B**) At 3, 5-hour post inoculation (hpi), the rate of conidial germination was measured and compared with the WT. (**C**) At 3, 5, 8, 12 and 20-hpi, the percentage of appressorium development was measured and compared with the WT. Results are the mean of three independent biological measurements. A total of 150 spores were counted for each replicate. Images were taken at each timepoint. Error bars denote SD. Asterisks indicate statistically significant differences (*p<0.05, **p<0.01, ***p<0.001, ****p<0.0001, two-way ANOVA with Tukey’s multiple comparisons test using GraphPad Prism 8). Scale bar is 10 μm.

**Figure 6.**
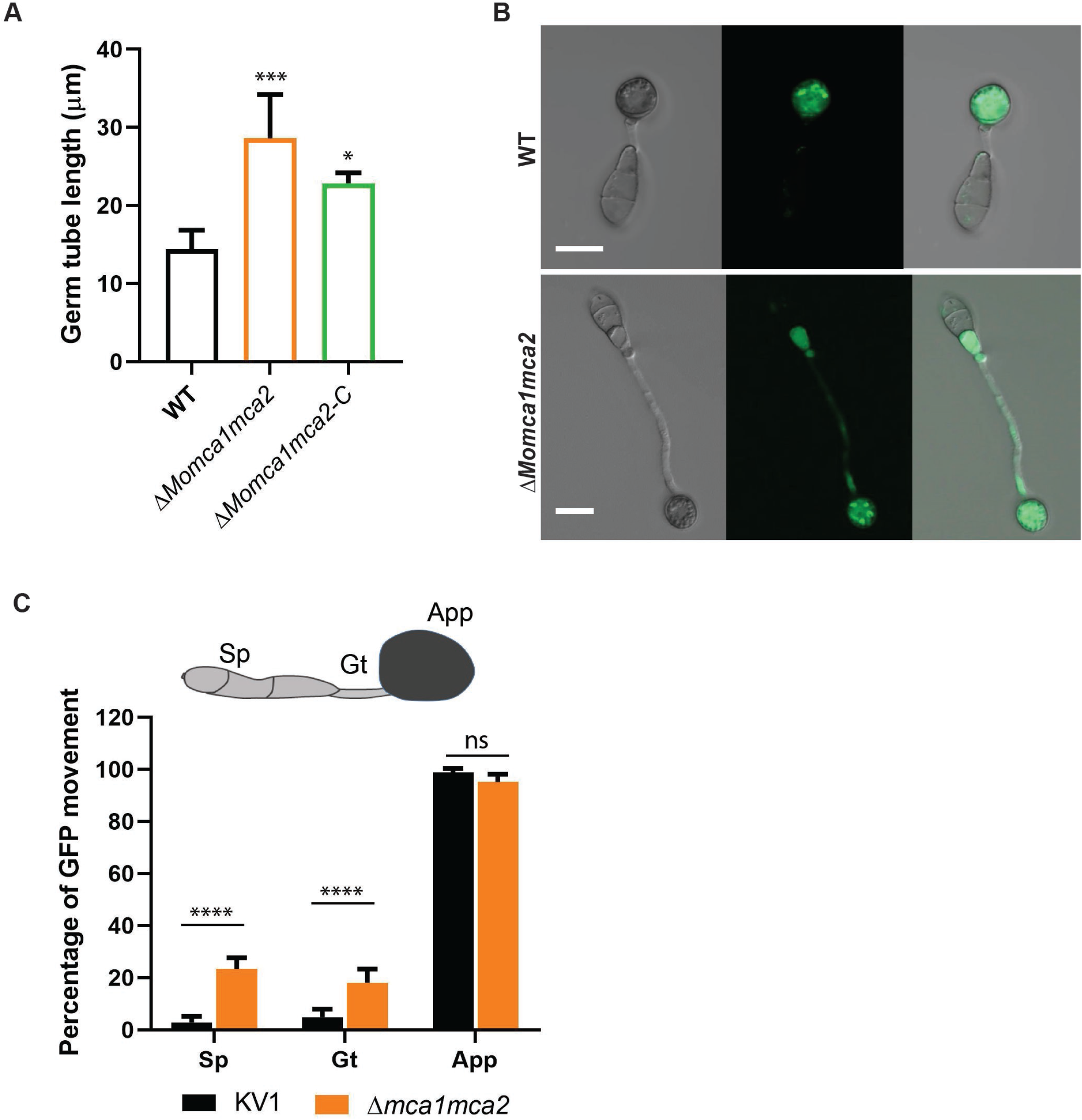
Δ *Momca1mca2* mutant strain developed long germ tubes during appressorium formation. (**A**) Average germ tube length of the WT, Δ*Momca1mca2* and Δ*Momca1mca2-C* mutant strains at 20 hours post incubation. (**B**)(**C**) Conidium content at 20h after incubation on inducible surface. Percentage of cytoplasmic eGFP movement into the appressorium in the Δ*Momca1mca2* compared to the WT. Results are the mean of three independent measurements. A total of 150 germinated spores with appressoria were counted for each replicate. Sp: spores, Gt: germ tube, App: appressorium. Error bars denote SD. ns: not significant. Asterisks indicate statistically significant differences (*p<0.05, ***p<0.001, ****p<0.0001, one-way ANOVA with Tukey’s multiple comparisons test using GraphPad Prism 8). Scale bar is 10 μm.

### *M. oryzae* metacaspases play a role in the clearance of protein aggregates

Based on previous studies implicating the role of Yca1 in maintaining protein homeostasis, we next investigated whether the mutants might alter protein stability during conidial germination and appressorium formation (7). In *M. oryzae*, the conidium contents are recycled into the appressorium, contributing to the next stages of appressorium development (20, 21). To observe the cytoplasmic protein content during appressorium formation, we used the KV1 strain that harbors a cytosolic eGFP reporter protein (24). Spores from WT KV1 and Δ*Momca1mca2* strains were collected and incubated on a hydrophobic surface for 20 hpi. Normally on hydrophobic surfaces, *M. oryzae* spores germinate and form a short germ tube with an immature appressorium within 6 hr. During this process, the conidial contents migrate into the incipient appressorium (20, 25). Using eGFP as a marker, we can observe the movement of proteins into the conidium during this developmental process. For the WT KVI strain, Fig. 6B and C show the expected cytoplasmic movement of proteins from the conidium into the appressorium. By contrast, in the Δ*Momca1mca2* strain, there is a delay of eGFP delivery due to the delay in the appressorium formation.

In yeast, Yca1 plays an essential role in protein quality control by promoting the removal of insoluble protein aggregates to maintain the fitness of new cells (7). Δ*yca1* mutant showed a significant increase of insoluble aggregates in the cells compared to the WT yeast strain (7). Therefore, we next asked if a similar phenotype could be observed in *M. oryzae*. We measured the insoluble protein aggregate fractions in *M. oryzae* WT, Δ*Momca1mca2* and the Δ*Momca1mca2-C* complemented strain under normal gel electrophoresis. We observed an increase of insoluble aggregates in the Δ*Momca1mca2* mutant compared to the WT and a restoration of the WT phenotype in the complemented strain (Fig. 7A; see also Fig. S3C and D). The observations support the hypothesis that *M. oryzae* metacaspases play an important role in preventing protein aggregates from accumulating during appressorium formation. Previous reports demonstrate that the Hsp70s aggregate-remodeling chaperones are overexpressed as a compensatory response to yeast Yca1 deletion (7). To further study the protein aggregation phenotype associated with metacaspase deletion, we assessed the expression of the 70-kDa heat shock protein, Hsp70, in the spores and mycelia lysates. Using western blot analysis, we detected the expression of Hsp70 in the soluble and insoluble fractions of WT, Δ*Momca1mca2* and Δ*Momca1mca2-C* strains (Fig. 7B). As figure 7 shows, Hsp70 was found to accumulate in the insoluble fraction of the Δ*Momca1mca2* mutant compared to the WT and complemented strains. Moreover, these results may lead to a better understanding of metacaspase roles in maintaining protein homeostasis in *M. oryzae* cells throughout its development and pathogenesis.

**Figure 7.**
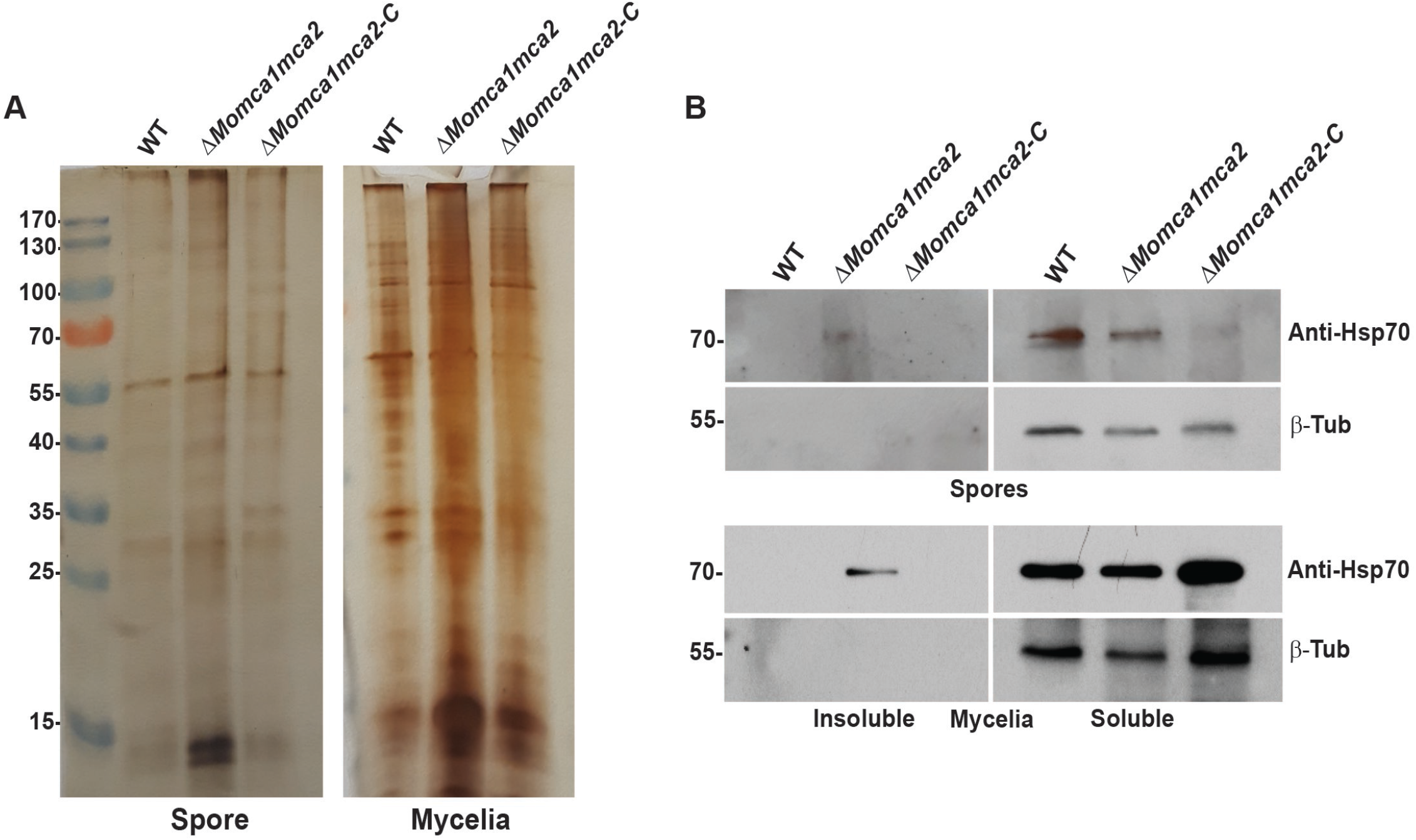
MoMca1 and MoMca2 play an essential role in clearance of insoluble aggregates. (**A**) Silver-stained 1D PAGE gel of insoluble fractions from equal amounts of total WT KV1, Δ*Momca1mca2* and Δ*Momca1mca2-C* spores and mycelia lysates. (**B**) Hsp70 accumulates in the insoluble fractions of Δ*Momca1mca2’s* spores and mycelia lysates. Western blot analysis of Hsp70 in the WT, Δ*Momca1mca2* and Δ*Momca1mca2-C* strains. Anti-HSP70 was used for protein detection in the insoluble fractions of spores and mycelia lysates. β-Tubulin was used as a loading control. The experiment was performed three times, and a representative blot is presented.

## DISCUSSION

Metacaspases are multifunctional proteases essential for normal physiology and pathology of non-metazoan species. These proteases have been extensively characterized in programmed cell-death and non-apoptotic processes. Although the metacaspase family exists in eukaryotic organisms, the functional roles of metacaspase proteins have been mostly uncharacterized in pathogenic fungi, especially plant pathogens. This prompted us to investigate the role of metacaspases in the rice blast fungus *M. oryzae*. In this study, we characterized the functions of MoMca1 and MoMca2 metacaspases in *M. oryzae*. Our studies found that *M. oryzae* metacaspases function not only in conidiation and germination, but also in the regulation of clearance of insoluble protein aggregates during normal fungal growth.

We bioinformatically identified two type 1-metacaspase proteins, MoMca1 and MoMca2, in the *M. oryzae* genome. These proteins contain an N-terminal prodomain rich in Q/N residues and a C-terminal peptidase C14 caspase domain. MoMca1 and MoMca2 adopt the structure of a caspase fold, similar to the yeast Yca1 core (Fig. S1A).

Due to the bioinformatic similarities of MoMca1 and MoMca2 with Yca1, we looked at the potential link in Ca^2+^ zymogen activation. In contrast with canonical caspases, metacaspases do not undergo dimerization for their activation. Instead, the activity of metacaspases depends on the presence of calcium ions (23, 26), with the only known exception being *Arabidopsis thaliana* AtMC9, whose activity was shown to be Ca^2+^-independent (5, 27). However, the *Arabidopsis* AtMCP2d metacaspase exhibits a strict Ca^2+^ dependence in millimolar concentration for its catalytic activation (Watanabe and lam, 2011). In Yca1, the addition of Ca^2+^activates the enzyme and leads to autoprocessing events (23). Our biochemical studies have identified MoMca1 and MoMca2 to be Ca^2+^-activated proteases, similar to the Yca1 metacaspase. First, we observed that the autocatalytic processing of MoMca1 and MoMca2 is stimulated by the presence of Ca^2+^ *in vitro*. In addition to the activation requirements for Ca^2+^, we found two cysteine residues that were essential for Yca1 activity were also important for MoMca1 and MoMca2 activity.

Similar to the role of canonical caspases, yeast Yca1 metacaspase positively regulates apoptosis under several stress conditions (28). In this report, we provide evidence that high level of oxidative stress inducers, such as H_2_O_2_ and menadione, upregulate the expression of *M. oryzae* metacaspase genes. These observations lead us to investigate the functional similarities between *M. oryzae* metacaspases and yeast Yca1. In early studies, Madeo and colleagues demonstrated that low concentrations of H_2_O_2_ facilitate the activation of Yca1 promoting yeast apoptosis (4). Lack of yeast Yca1 activity confers resistance to oxidative stress conditions and a higher survival rate under them (4). Here, we provide evidence that both MoMca1 and MoMca2 can substitute the Yca1 function in mediating oxidative stress-induced apoptosis in yeast. Although MoMca1 and MoMca2 functionally complement the oxidative stress phenotype in yeast, whether the *M. oryzae* metacaspases respond to similar external stimuli remains unclear. Moreover, Δ*Momca1mca2* strain shows resistance to H_2_O_2_ and menadione similar to Δ*yca1*.

Additionally, the deletion of either *MoMca1* or *MoMca2* in *M. oryzae* impairs developmental processes, while having no significant reduction in vegetative growth rate on solid media. We demonstrate that both conidiation and symptom development are metacaspase-dependent processes in *M. oryzae*, suggesting a role for these proteins in cell development. The Δ*Momca1mca2* strain was significantly impaired in proper conidiation and pathogenicity.

To be a successful pathogen, *M. oryzae* undergoes several morphological changes to colonize and proliferate inside the rice cells (19). Once the conidium attaches to the leaf cuticle, it germinates and develops a germ tube that will lead to the formation of the melanized appressorium. This morphogenesis process is tightly regulated by the cell cycle checkpoints and occurs within hours (2-24h) after conidium attachment (29). Surprisingly, we discovered that Δ*Momca1mca2* mutant exhibits delayed conidial germination under inducible conditions, suggesting that these proteins contribute to initial processes that precede appressorium morphogenesis. This delayed phenotype in the Δ*Momca1mca2* mutant disrupts the timing of the appressorium maturation process. Moreover, we observed a delay in the movement of conidial contents into the appressorium. It is well known that during appressorium development, the conidial contents are recycled and delivered to the appressorium leading to conidial degradation (25). This multistage process is highly regulated by cell cycle progression and autophagy-dependent cell death (25, 29, 30). However, how *M. oryzae* metacaspases crosstalk with autophagy processes to regulate appressorium development remains largely unknown.

In other systems, metacaspases display varied roles in pathogenicity and oxidative stress resistance. For instance, the filamentous fungus, *Podospora anserina*, contains two metacaspases and the deletion of both metacaspase genes leads to lower growth rate and fertility indicating a function in developmental processes (15). However, a single deletion strain, *PaMca1* has a lifespan-prolonging effect and resistance response to oxidative stress (15). By contrast, the CasA/CasB metacaspases in the human pathogen *Aspergillus fumigatus* showed no resistance to oxidative stress or to other apoptosis-induced agents. However, Δ*casA*Δ*casB* mutant displays a growth defect under ER stress conditions, indicating that these proteases confer vital cellular functions under certain stress conditions rather than an involvement in cell death processes (14). Recent studies identified a single metacaspase gene in *Ustilago maydis* that exhibits impaired growth under either normal growth or oxidative stress conditions (17). The *U. maydis mca1* mutant also displayed a reduction in pathogenicity when compared to the wild-type, demonstrating that the Mca1 in *U. maydis* is required for cellular homeostasis during cell development (17). In this study, we found that *M. oryzae* Δ*Momca1mca2* mutant exhibited a reduction in pathogenicity on rice plants. With these observations, we can infer that metacaspase functions vary among different organisms, and have partial redundancy or antagonistic functions under various settings.

In *S. cereviciae,* Yca1 metacaspase appears to be critical for the removal of insoluble protein aggregates during physiological growth conditions. The deletion of *yca1* was associated with an accumulation of insoluble aggregates during logarithmic growth that correlated with an enrichment on vacuolization and stress-response chaperones (7). Due to previous findings about Yca1 promoting clearance of proteins aggregates, we decided to investigate the involvement of our metacaspases on this process. It is noteworthy that in Δ*Momca1mca2* mutant we observed the accumulation of insoluble protein aggregates during normal growth conditions not present in the control. These findings clearly suggest that *M. oryzae* metacaspases contribute to the removal of insoluble aggregates to maintain optimal fitness during fungal growth. However, we are not clear how *M. oryzae* mediates the clearance of insoluble aggregates during physiological growth. We initially hypothesized that the accumulation of the insoluble aggregates may have a direct effect in delaying conidial germination in *M. oryzae*. Though, a recent study on protein aggregates in *M. oryzae* found no effect on germination when inducing protein aggregates in the spores (31). Although protein aggregates were present in large quantities in catalytic mutants, these spores did not have delayed germination (31). This data suggests that germination itself in the Δ*Momca1mca2* mutant is affected in other ways, independent of protein aggregate accumulation, potentially through some other metacaspase regulatory function. The discovery of metacaspase substrates will give us new insight into the mechanism of activation *in vivo* and the downstream events.

Our work characterizing the role of the *M. oryzae* metacaspases, MoMca1 and MoMca2, has demonstrated that biochemically there are several shared properties with the Yca1 metacaspase. MoMca1 and MoMca2 are Ca^2+^ dependent peptidases. While MoMca1 and MoMca2 are dispensable during vegetative growth, they are critical for inducing apoptosis under oxidative stress conditions. More importantly, the deletion of these genes leads to an accumulation of protein aggregates. The most striking phenotype of the *M. oryzae* metacaspases is that they are critical for sporulation and subsequently pathogenicity in rice blast disease progression.

## MATERIALS AND METHODS

### Strains and culture conditions

The *M. oryzae* strain KV1 was used as the wild-type strain throughout this research (24); Table S1). This strain and its transformants were cultured on complete media (CM; 1% (W/V) glucose, 0.2% (W/V) peptone, 0.1% (W/V) yeast extract and 0.1% (W/V) casamino acids, 0.1% trace elements, 0.1% vitamin solution and 1X nitrate salts) and incubated at 25°C under 12 hrs light/dark cycles for 5–12 days. For DNA and RNA extraction, all strains were grown on CM liquid media at 25°C with agitation for 48h. For sporulation rates, strains were grown on at least three independent CM plates. After 12 days of growth, spores were harvested and counted using a hemocytometer (Corning). To observe vegetative mycelial growth under stress, 10mM H_2_O_2_ and 150μM menadione were individually added to CM agar medium. A mycelial disc (3.5 mm in diameter) of each strain was inoculated on stress media containing either H_2_O_2_ or menadione, and the growth rate was assessed by measuring culture diameters after 5 days of growth (unless otherwise stated). Plate images were taken with Epson perfection V700 photo scanner. For routine cloning, the *Escherichia coli* DH5α strain was grown in 2xYT broth at 37°C. The *Saccharomyces cerevisiae* yeast strain BY4741 was grown in Synthetic Dropout (SD) media as described (32) at 30°C.

### Generation of Plasmids

The *M. oryzae Mca1 and Mca2* coding sequences were amplified by PCR using *M. oryzae* cDNA as a template. For *in vitro* transcription/translation reactions, *S. cerevisae Yca1, M. oryzae Mca1 and Mca2* coding sequences were cloned into pET15b bacterial expression plasmid containing an N-terminal 6XHis tag. Single amino acid mutations were introduced via Quick Change site directed mutagenesis.

For yeast expression, *MoMca1* and *MoMca2* full length and variants were cloned into the XhoI/HindIII sites on a galactose-driven yeast expression vector pESC-Leu (kind gift from Vincent S. Tagliabracci).

To test the redundancy of *M. oryzae* Mca1 and Mca2 to yeast metacaspase, Yca1, we obtained an Yca1 knockout yeast strain (*yca1*Δ::KANMX; Dharmacon) derived from BY4741 strain (33).

### Sequence analysis

The *MoMca1* and *MoMca2* coding gene sequences were obtained from the *Magnaporthe oryzae* database http://fungi.ensembl.org/Magnaporthe_oryzae/Info/Index. Metacaspase sequences were collected using the MoMca1 (A4QTY2.2) sequence as a query for PSI-BLAST (34) (5 iterations, E-value cutoff 0.005) against a sequence database containing the RefSeq representative prokaryotic genome set (1684 genomes, from Aug 28, 2018) and latest representative eukaryotic genomes (from Aug 28, 2018). Collected sequences (2712) were clustered using CLANS (35) and colored according to taxonomy. A multiple sequence alignment of representative sequences was generated using MAFFT (36) and colored according to conservation: mainly hydrophobic (yellow), mainly small (gray), active site (black), and calcium activation (red). To identify a structure template for MoMca1, the sequence (A4QTY2.2) was submitted to the HHPRED server (37) to search against the PDB database, which confidently identified the Yca1 structure (4f6o) as a top template (probability 100%, score 2.2e-38). The HHPRED pairwise alignment between MoMca1 and Yca1 was used to generate a structure model for MoMca1 using SwissModel (38).

### Fungal transformation and complementation

Targeted gene replacements for *MoMca1* and *MoMca2* were carried out using the PCR-based split marker method (39) in which the *HPH* gene (1.4-kb), conferring resistance to hygromycin, and *ILV1* gene (2.8-kb), conferring resistance to sulfonylurea, replaced the coding sequence of each gene. Approximately 1 kb upstream and downstream of the gene were used for homologous recombination. The flanking sequences (~1kb) of the *MoMca1* and *MoMca2* genes were amplified and fused to the selectable marker by PCR (Figure S1A). PCR products were transformed into *M. oryzae* WT protoplasts. Positive transformants carrying homologous gene replacement of the gene of interest were confirmed by PCR using the nested primers shown in Table S2.

To generate the complemented strain, the full length of *MoMca1* and *MoMca2* genes including their native promoter were amplified and inserted into pBGt (40) fungal expression vector by using the Gibson Assembly approach (NEB BioLabs). The pBGt-MoMca1Mca2 expression vector was transformed into Δ*Momca1mca2* protoplasts and selected on gentamicin-containing plates. The complementation strain was identified by PCR (Table S2). Single-spores isolation was performed for all the strains.

### Spore germination and appressorium development assays

Spores were harvested from 10-day-old cultures, filtered through two layers of Miracloth (Millipore Sigma), and re-suspended to a concentration of 5×10^4^ spores/ml in sterile water. For conidial germination, 20μl of conidial suspension were placed on glass cover slips (Fisher Scientific, St Louis, MO, USA) and incubated at 24°C in a humid chamber. The percentage of conidial germination and appressorium formation were assessed at 3, 5, 8,12 and 20 hours post incubation. Average mean values were determined from 150 conidia and performed in three independently experiments. Germ tube lengths were measured using ImageJ. All imaging was performed on a Zeiss LSM 800 confocal microscope and images were converted using ImageJ (NIH).

### Pathogenicity assays

Spores from WT and transformants were harvested from 10-day-old cultures on 0.2% gelatin as previously described (41). Three-week old rice seedlings (YT16) were spray-inoculated with 2×10^4^ spores/ml conidial suspension. Inoculated plants were placed in a humidity chamber for 24 h, and then transferred back to the growth chamber with 12 hrs light/dark cycles. Disease severity was assessed at 7-day post inoculation. Photographs of diseased rice leaves were taken. The number of pixels under lesion areas and healthy areas of diseased leaves were calculated by using ImageJ.

### Yeast transformation

Yeast transformations were performed using the Lithium acetate (LiAc) method as described in (42). Briefly, yeast cells were grown overnight at 30°C in YPD media. The overnight cultures were diluted in 50ml of fresh YPD (adjusted to an OD_600_ of 0.1) and then cultured until reaching an OD of 0.5-0.7. After incubation, the yeast cells were collected by centrifugation and washed twice with solution 1 (1X TE, 1M LiAC, pH 7.5 with acetic acid). The pellet was resuspended with 500μl of solution 1. In 100μl of yeast cells, 1ug of DNA, 5μl DNA carrier, and 5μl of 100% DMSO were added and gently mixed. 700μl of solution 2 (solution 1 supplemented with 50% PEG) was added and incubated at 30°C for 30 minutes. The yeast cells were heat shocked at 42°C for 15 min, and washed with TE pH 7.5. Samples were plated on Synthetic drop-out (SD) media minus Leucine or Uracil and incubated at 30°C for 2-3 days.

### Survival Assay

Yeast survival assay was performed as described in (4). Briefly, yeast strains harboring the expression vector were grown overnight in 3ml liquid SD-Leu glucose media at 30°C. The overnight cultures were diluted in 10ml of fresh media (adjusted to an OD_600_ of 0.05) and then cultured until reaching an OD_600_ of 0.4-0.6. Then, the yeast cells were collected and resuspended in SD-Leu supplemented with 2% galactose and 1% raffinose. Cell counts were equalized based on OD_600_ before treatment. For stimulation of yeast apoptosis, cells were treated with the final concentration of 1.2mM H_2_O_2_ and incubated for 24h at 30°C. An aliquot of each culture was 10-fold diluted in sterile water and plated in SD glucose plates. Colonies were counted after 2-3 days and treated samples were compared with non-treated samples to calculate survival rate. This experiment was conducted in biological triplicates.

### RNA extraction, cDNA synthesis and qRT-PCR analysis

To quantify metacaspase expression under oxidative stress conditions, the fungal strains were grown on liquid CM for 48h at 25°C with agitation (200 rpm) before treatment with or without 10mM H_2_O_2_ and 150μM menadione for 2h. Mycelia were harvested and ground with a pestle and mortar in liquid nitrogen, and stored at −80°C. A total of 100mg of ground mycelia was used to performed RNA extractions. Total RNA was isolated using the RNeasy Plant mini plus kit from Qiagen. RNA concentration was measured via NanoDrop, and cDNA was generated using the iScript cDNA Synthesis Kit (Quanta). Primers were designed to amplify 100 to 150 base pairs of each target gene (Table S2) and tested for efficiency. Transcripts were quantified on a CFX384 Touch Real-Time PCR Detection System using the iTaq Universal SYBR Green Supermix (Quanta) and 500nM primers. Relative gene expression for each target gene was calculated by the −ΔΔCt method. The expression of each gene was normalized against the *M. oryzae* actin gene (*ACT1*). Results are given as the average of three technical replications and three biological replications.

### In vitro transcription/translation and purification

Coupled transcription/translation reactions were carried out per manufacturer’s instructions (Promega, L1170). Briefly, rabbit reticulocyte lysate was incubated with 1μg of the appropriate expression plasmid and 2mL of [^35^S] L-methionine (1,000 Ci/mMol at 10mCi/mL) (Perkin Elmer, NEG709A) for 90 minutes at room temperature. Reactions were diluted in 1mL of 50mM Tris-HCl (pH=7.5), 10mM NaCl, and 1mM DTT. 100μL of DEAE-Sepharose (Millipore Sigma, DFF100) slurry was added to the reactions and incubated on a nutator for 30 min at room temperature. The slurry was washed three times with 1mL of 50mM Tris-HCl (pH=7.5), 10mM NaCl, and 1mM DTT, and eluted with 500μL of 50mM Tris-HCl (pH=7.5), 500mM NaCl, and 1mM DTT. Eluents were collected and incubated with 50μL of packed Ni-NTA resin (Bio-Rad His-Pur 88222) at room temperature with nutating for 30 min. The beads were washed three times with 1mL of 50mM Tris-HCl (pH=7.5), 150mM NaCl, 20mM imidazole and 1mM DTT. Proteins were eluted using 100μL of 50mM Tris-HCl (pH=7.5), 150mM NaCl, 300mM imidazole and 1mM DTT.

### Protease assays

Protease reactions were carried out in a buffer solution containing 50mM Tris-HCl (pH 7.5), 150mM NaCl, 1mM DTT and with or without 1mM CaCl_2_. Reactions were incubated at 30°C overnight and stopped by the addition of Laemmli buffer containing β-mercaptoethanol and boiling. Reaction products were resolved on a 12% polyacrylamide gel by electrophoresis and autoradiography.

### Protein aggregates assays

Protein fractions from spores and mycelial cell lysates were collected as described by (7) with some modifications. Briefly, spores were harvested from 10-day-old cultures, filtered through two layers of Miracloth (Milipore Sigma), and re-suspended to a concentration of 5×10^5^ spores/ml in sterile water. Spores were recovered by centrifugation and pellets were frozen down and stored at −80°C for later processing. Mycelium was harvested and ground with a pestle and mortar in liquid nitrogen, and stored at −80°C. A total of 100mg of ground mycelia was used to performed protein extractions. Pellet samples were thawed in equal volumes of lysis buffer (0.1% Triton X-100, 50mM Tris pH 7.4, 1mM EDTA, and 1% glycerol supplemented with 5mM Na3VO4 and 1mM PMSF) and acid-washed glass beads (Sigma-Aldrich). Samples were vortexed for 20 minutes; 1min ON/OFF cycles. To remove the cell debris, samples were centrifuged at 2000 rpm. The total cell lysate was collected and protein concentration was measured and normalized. The cell lysate was centrifuged at 15,000 x g for 15 min at 4°C to separate the soluble (supernatant) and insoluble (pellet) fractions. Pellets were washed twice with lysis buffer containing 10% Triton X-100. Pellets were treated with 4M Urea and boiled in 2x Laemmli sample buffer for 5 minutes. Proteins were loaded on 12.5% SDS–polyacrylamide gel electrophoresis for 1D PAGE separation for silver staining or transferred to polyvinylidene difluoride membranes. HSP70 expression was detected with mouse anti-HSP70 (1:5,000; Abcam) antibody and β-Tubulin (polyclonal Anti-β-TUB; Santa Cruz) was used as a loading control.

### Statistical analysis

All data are given as mean ± standard deviation from at least three independent experiments (unless stated otherwise). Each experiment was conducted in triplicate. Statistical analyses were performed by using one-way ANOVA analysis and Tukey’s multiple comparisons test. A *p* value of < 0.05 was considered significant.

## ACKNOWLEDGEMENTS

We thank members of the Orth lab for their discussions and editing and Amanda Casey for sharing technical expertise on yeast systems. This work was funded by NIH grants R01 GM115188 and RO1 GM113079, Once Upon a Time…Foundation and the Welch Foundation I-1561. Dr. Kim Orth is a Burroughs Welcome Investigator, a Beckman Young Investigator, a W. W. Caruth, Jr. Biomedical Scholar, and the Earl A. Forsythe Chair in Biomedical Science.

## Author contribution

J.F. and K.O. conceived the project and designed the experiments. J.F., V.L., H.G., L.K., N.G., and M.P. conducted the experiments. J.F. and K.O. wrote the manuscript with input from all authors. K.O. supervised the project.

---

**Supplemental figure 1.** (**A**) The structure (PDB: 4f6o) is colored the same as the alignment, with functional residues shown in stick. The cleavage site is disordered, shown by a dashed line (magenta). The calcium is represented by a samarium ion from a superimposed homolog (PDB: 4afp). (**B**) Split marker method diagram. SP stands for selectable primer. *ILV1* and *HPH* genes that confer resistance against sulfonylurea drug and hygromycin selection, respectively. (**C**) Deleted mutant stains were confirmed by PCR. Primers P1 and P4 amplified 1kb upstream and downstream for the *MoMca1* and *MoMca2* genes. Top gels: Left (*MoMca1*); Right: (*MoMca2*). Primers P5 and P6 amplified internal sequences of *Momca1* and *Momca2* genes. Metacaspase genes in the WT genome contain several introns. Bottom gels: Left (*MoMca1*); Right: (*MoMca2*). *Momca1* gene was replaced in Δ*Momca1mca2* mutant with *hph* gene that confers resistance to hygromycin selection. The complemented strain, Δ*Momca1mca2-C,* was generated by cloning the *MoMca1* and *MoMca2* ORFs with their respective 1kb native promoter into the binary pBHt2 vector and transfected into Δ*Momca1mca2* protoplasts.

**Supplemental figure 2.** (**A**)(**B**) Radial growth on complete media (CM) was not impaired in mutant strains carrying gene deletions in *MoMca1, MoMca2* or in the double mutant. KV1 is the wild-type isolate (WT) used in this study. Images were taken at 5 days after incubation. (**C**) Sporulation was impaired in strains lacking functional *Mca* genes, but more significantly reduced in Δ*mca1mca2* strain during growth on CM. Values represent the mean of three independent replicates. Error bars denote SD. Asterisks indicate statistically significant results: \*\*\**P*<0.001 (one-way ANOVA with Tukey’s multiple comparisons test using GraphPad Prism 8).

**Supplemental figure 3.** Metacaspase mutants Δ *Momca1,* ΔMo*mca2* and Δ*Momca1mca2* are not sensitive to the osmolytes 1M Sorbitol and 1M NaCl and cell wall assembly inhibitor Congo Red. (**A**) Stressors were added to CM at the concentrations shown. Images were taken after 5 days of growth. (**B**) Relative density of the aggregates on insoluble fractions in the spores and mycelia lysates. The relative density was measured using ImageJ. The relative values were obtained from three independent experiments with technical repetitions. Error bars denote SEM. Asterisks indicate statistically significant differences (\*\**p*<0.01, \*\*\*\**P*<0.0001, one-way ANOVA with Tukey’s multiple comparisons test using GraphPad Prism 8).

**Table S1.** Strains used in this study

**Table S2**. Oligonucleotide primers used in this study.

## REFERENCES

1. Shrestha A, Megeney L. 2012. The non-death role of metacaspase proteases. Frontiers in Oncology 2:78.

2. Tsiatsiani L, Van Breusegem F, Gallois P, Zavialov A, Lam E, Bozhkov PV. 2011. Metacaspases. Cell Death And Differentiation 18:1279.

3. Carmona-Gutierrez D, Fröhlich KU, Kroemer G, Madeo F. 2010. Metacaspases are caspases. Doubt no more. Cell Death And Differentiation 17:377.

4. Madeo F, Herker E, Maldener C, Wissing S, Lächelt S, Herlan M, Fehr M, Lauber K, Sigrist SJ, Wesselborg S, Fröhlich K-U. 2002. A caspase-related protease regulates apoptosis in yeast. Molecular Cell 9:911–917.

5. Vercammen D, van de Cotte B, De Jaeger G, Eeckhout D, Casteels P, Vandepoele K, Vandenberghe I, Van Beeumen J, Inze D, Van Breusegem F. 2004. Type II metacaspases Atmc4 and Atmc9 of *Arabidopsis thaliana* cleave substrates after arginine and lysine. J Biol Chem 279:45329–36.

6. Tsiatsiani L, Van Breusegem F, Gallois P, Zavialov A, Lam E, Bozhkov PV. 2011. Metacaspases. Cell death and differentiation 18:1279–1288.

7. Lee REC, Brunette S, Puente LG, Megeney LA. 2010. Metacaspase Yca1 is required for clearance of insoluble protein aggregates. Proceedings of the National Academy of Sciences 107:13348–13353.

8. Lee REC, Puente LG, Kærn M, Megeney LA. 2008. A non-death role of the yeast metacaspase: Yca1p alters cell cycle dynamics. PLOS ONE 3:e2956.

9. Khan MAS, Chock PB, Stadtman ER. 2005. Knockout of caspase-like gene, *YCA1*, abrogates apoptosis and elevates oxidized proteins in *Saccharomyces cerevisiae*. Proceedings of the National Academy of Sciences of the United States of America 102:17326–17331.

10. Guaragnella N, Pereira C, Sousa MJ, Antonacci L, Passarella S, Côrte-Real M, Marra E, Giannattasio S. 2006. YCA1 participates in the acetic acid induced yeast programmed cell death also in a manner unrelated to its caspase-like activity. FEBS Letters 580:6880–6884.

11. Guaragnella N, Bobba A, Passarella S, Marra E, Giannattasio S. 2010. Yeast acetic acid-induced programmed cell death can occur without cytochrome c release which requires metacaspase YCA1. FEBS Letters 584:224–228.

12. Guaragnella N, Passarella S, Marra E, Giannattasio S. 2010. Knock-out of metacaspase and/or cytochrome c results in the activation of a ROS-independent acetic acid-induced programmed cell death pathway in yeast. FEBS Letters 584:3655–3660.

13. Shrestha A, Brunette S, Stanford WL, Megeney LA. 2019. The metacaspase Yca1 maintains proteostasis through multiple interactions with the ubiquitin system. Cell Discovery 5:6.

14. Richie DL, Miley MD, Bhabhra R, Robson GD, Rhodes JC, Askew DS. 2007. The *Aspergillus fumigatus* metacaspases CasA and CasB facilitate growth under conditions of endoplasmic reticulum stress. Mol Microbiol 63:591–604.

15. Hamann A, Brust D, Osiewacz HD. 2007. Deletion of putative apoptosis factors leads to lifespan extension in the fungal ageing model *Podospora anserina*. Molecular Microbiology 65:948–958.

16. Cao Y, Huang S, Dai B, Zhu Z, Lu H, Dong L, Cao Y, Wang Y, Gao P, Chai Y, Jiang Y. 2009. *Candida albican*s cells lacking CaMCA1-encoded metacaspase show resistance to oxidative stress-induced death and change in energy metabolism. Fungal Genetics and Biology 46:183–189.

17. Mukherjee D, Gupta S, Saran N, Datta R, Ghosh A. 2017. Induction of apoptosis-like cell death and clearance of stress-induced intracellular protein aggregates: dual roles for *Ustilago maydis* metacaspase Mca1. Mol Microbiol 106:815–831.

18. Wang X, Wang Y, Zhou Y, Wei X. 2014. Farnesol induces apoptosis-like cell death in the pathogenic fungus *Aspergillus flavus*. Mycologia 106:881–888.

19. Fernandez J, Orth K. 2018. Rise of a cereal killer: The biology of *Magnaporthe oryzae* biotrophic growth. Trends Microbiol 26:582–597.

20. Ryder LS, Talbot NJ. 2015. Regulation of appressorium development in pathogenic fungi. Current opinion in plant biology 26:8–13.

21. Talbot NJ. 2019. Appressoria. Curr Biol 29:R144–R146.

22. de Jong JC, McCormack BJ, Smirnoff N, Talbot NJ. 1997. Glycerol generates turgor in rice blast. Nature 389:244–244.

23. Wong AH, Yan C, Shi Y. 2012. Crystal structure of the yeast metacaspase Yca1. J Biol Chem 287:29251–9.

24. Kankanala P, Czymmek K, Valent B. 2007. Roles for rice membrane dynamics and plasmodesmata during biotrophic invasion by the blast fungus. The Plant cell 19:706–724.

25. Veneault-Fourrey C, Barooah M, Egan M, Wakley G, Talbot NJ. 2006. Autophagic fungal cell death is necessary for infection by the rice blast fungus. Science 312:580.

26. Moss CX, Westrop GD, Juliano L, Coombs GH, Mottram JC. 2007. Metacaspase 2 of *Trypanosoma brucei* is a calcium-dependent cysteine peptidase active without processing. FEBS Letters 581:5635–5639.

27. Watanabe N, Lam E. 2011. Calcium-dependent activation and autolysis of arabidopsis metacaspase 2d. Journal of Biological Chemistry 286:10027–10040.

28. Madeo F, Carmona-Gutierrez D, Ring J, Büttner S, Eisenberg T, Kroemer G. 2009. Caspase-dependent and caspase-independent cell death pathways in yeast. Biochemical and Biophysical Research Communications 382:227–231.

29. Saunders DGO, Aves SJ, Talbot NJ. 2010. Cell cycle–mediated regulation of plant infection by the rice blast fungus. The Plant Cell 22:497–507.

30. Osés-Ruiz M, Sakulkoo W, Littlejohn GR, Martin-Urdiroz M, Talbot NJ. 2017. Two independent S-phase checkpoints regulate appressorium-mediated plant infection by the rice blast fungus *Magnaporthe oryzae*. Proceedings of the National Academy of Sciences 114:E237–E244.

31. Rogers AM, Egan MJ. 2020. Autophagy machinery promotes the chaperone-mediated formation and compartmentalization of protein aggregates during appressorium development by the rice blast fungus. Molecular Biology of the Cell 31:2298–2305.

32. Salomon D, Sessa G. 2010. Identification of growth inhibition phenotypes induced by expression of bacterial type III effectors in yeast. J Vis Exp:(37):1865.

33. Baker Brachmann C, Davies A, Cost GJ, Caputo E, Li J, Hieter P, Boeke JD. 1998. Designer deletion strains derived from *Saccharomyces cerevisiae* S288C: A useful set of strains and plasmids for PCR-mediated gene disruption and other applications. Yeast 14:115–132.

34. Altschul SF, Madden TL, Schaffer AA, Zhang J, Zhang Z, Miller W, Lipman DJ. 1997. Gapped BLAST and PSI-BLAST: a new generation of protein database search programs. Nucleic Acids Res 25:3389–402.

35. Frickey T, Lupas A. 2004. CLANS: a Java application for visualizing protein families based on pairwise similarity. Bioinformatics 20:3702–4.

36. Katoh K, Standley DM. 2014. MAFFT: iterative refinement and additional methods. Methods Mol Biol 1079:131–46.

37. Soding J, Biegert A, Lupas AN. 2005. The HHpred interactive server for protein homology detection and structure prediction. Nucleic Acids Res 33:W244–8.

38. Arnold K, Bordoli L, Kopp J, Schwede T. 2006. The SWISS-MODEL workspace: a web-based environment for protein structure homology modelling. Bioinformatics 22:195–201.

39. Goswami RS. 2012. Targeted gene replacement in fungi using a split-marker approach. Methods Mol Biol 835:255–69.

40. Kim HS, Park SY, Lee S, Adams EL, Czymmek K, Kang S. 2011. Loss of cAMP-dependent protein kinase A affects multiple traits important for root pathogenesis by *Fusarium oxysporum*. Mol Plant Microbe Interact 24:719–32.

41. Fernandez J, Marroquin-Guzman M, Wilson RA. 2014. Evidence for a transketolase-mediated metabolic checkpoint governing biotrophic growth in rice cells by the blast fungus *Magnaporthe oryzae*. PLoS Pathog 10:e1004354.

42. Salomon D, Sessa G. 2010. Identification of growth inhibition phenotypes induced by expression of bacterial type III effectors in yeast. J Vis Exp.

